# Development of a multi-species luciferase-based double antigen ELISA for the detection of antibodies against Influenza A virus H5 clade 2.3.4.4b

**DOI:** 10.64898/2026.01.05.697617

**Authors:** Andrea Aebischer, Anne Günther, Ronja Piesche, Kerstin Wernike, Magdalena Murr, Timm Harder, Donata Hoffmann, Christian Grund, Beatriz Bellido Martín, Ron Fouchier, Christoph Kreer, Katharina Daniel, Leon Ullrich, Rosanne Sprute, Marco Roller, Lukas Reese, Josefine Wassermann, Gereon Schares, Maryna Kuryshko, Elsayed M. Abdelwhab, Martin Beer

## Abstract

The highly pathogenic avian influenza viruses (HPAIV) of subtype H5N1 represent a major threat to animal and public health. The current panzootic with H5 clade 2.3.4.4b has caused numerous, widespread outbreaks in various domestic and wild avian species with high mortalities, massive losses and a high frequency of spillover events to unexpected novel mammalian hosts such as dairy cows. The global H5N1 situation raises serious concerns about zoonotic risks due to effective mammal-to-mammal transmission. Therefore, it is critical to increase surveillance intensity of a broadened species range, particularly at the human-animal interface. For this purpose, reliable and cost-effective serological tools that are easy to perform and suitable for high-throughput screenings are critically needed. The newly developed double antigen ELISA format employing a luminescence-based detection technology has demonstrated to comply with such prerequisites. The assay allowed the detection of H5-specific antibodies even early after infection or vaccination in a wide range of birds and mammals including humans. It further demonstrated superior analytical sensitivity and high specificity for antibodies directed against H5 hemagglutinin of clade 2.3.4.4b as no cross-reactivity with other hemagglutinin subtypes was observed. Thus, the assay represents a valuable contribution to existing serological diagnostic tests for a clade-optimized detection of influenza A virus antibodies in a broad range of species.

**Importance:** The ongoing HPAIV H5N1 panzootic has caused numerous outbreaks in domestic and wild animals with frequent spillover events to unexpected host species, which underscores the importance of an intensified surveillance. However, sensitive and specific multi-species serological assays represent a major gap. For this purpose, we developed a novel double antigen ELISA which employs an innovative luminescence-based read-out strategy. The test allowed a highly sensitive and specific detection of H5-specific antibodies even early after infection or vaccination in a wide range of avian and mammalian species including humans. It therefore represents a significant contribution to improving species-independent serological diagnostic tools for the detection of influenza A virus antibodies.

## Introduction

Infections with highly pathogenic avian influenza (HPAI) viruses have severe consequences for animal and human health as currently demonstrated by the global spread of the HPAI H5N1 clade 2.3.4.4b. Since 2016 this clade caused outbreaks with devastating economic losses in domestic poultry as well as previously unseen massive die-offs in different wild bird species (1). Furthermore, frequent spillovers in more than 80 mammalian species have been reported and associated with high morbidities and mortalities (2–4) which raises serious concerns about the threat to endangered species and biodiversity (5, 6). The incursion of H5N1 to Northern and Southern America not only led to high numbers of fatalities among marine mammals, but also revealed evidence of a direct transmission between the animals (7–10). Mammalian transmission was previously also suspected during outbreaks in fur farms in Spain and Finland (11, 12). In 2024, H5N1 clade 2.3.4.4b was detected for the first time in dairy cattle in the United States (U.S.) and spread rapidly to affect more than 1000 dairy farms in 18 U.S. states since that time (13). Subsequently, deadly infections of cats and other mammals in contact with highly infectious raw milk of affected farms as well as of farm workers in close contact to infected poultry or cattle were reported (14, 15). To date, sustained person-to-person transmission has not been documented but the evidence of mammalian adaptations in the viruses isolated from different species proved the propensity of the virus to evolve towards more efficient replication in mammals including humans (2, 16). All these studies emphasize the necessity of increasing surveillance intensity of a broadened spectrum of potential host species, especially at the human-animal interface. For this purpose, reliable multi-species serological tools are urgently needed (17). Enzyme-linked immunosorbent assays (ELISA) represent one of the most appropriate test systems since they are easy to conduct and suitable for automation allowing a rapid and cost-effective analysis of large numbers of samples.

Over the last few years, several ELISAs for the detection of antibodies against hemagglutinin (HA) subtype H5 have been developed using different approaches. Even though, antibody-competition formats allow a species-independent application, most assays have primarily been developed and validated for galliform poultry. Their performance in other domestic and wild avian species often showed diverging results and comparative data for mammalian species is mostly not available (18–24). Though innovative, other detection strategies still rely on species- or antibody isotype-dependent reagents (25–29).

In order to enable a clade-optimized, specific detection of anti-H5 antibodies which is completely independent of species and isotype, we developed a novel assay based on the previously described luciferase immunoprecipitation system (LIPS) technology (30, 31). The principle of the liquid-phase double antigen ELISA is based on the capture of antibodies by their simultaneous binding to an immobilized capture antigen and a nanoluciferase-labeled detector antigen. The reporter gene produces a high intensity, glow-type luminescence which allows quantitation of antibody titers in small sample volumes with low background auto-luminescence (32). In addition, the system does not require a purification of the detector antigen which facilitates production and reduces costs. Our data show that the novel assay combined the outstanding sensitivity of the LIPS technique with a fast turnaround time and the capability for a robust high-throughput application of a standard solid-phase ELISA. Thus, the format represents a valuable and versatile contribution to existing H5-specific serological tests to improve antibody detection in a broad range of host species.

## Material and Methods

### Cloning

The full-length H5 hemagglutinin (HA) protein comprising amino acids (aa) 17 – 530 from influenza virus isolate A/arctic tern/Germany-SH/AI03798/2022 (A/H5N1, clade 2.3.4.4b) (EPI2242291) was amplified from a codon-optimized synthetic gene (GeneArt, Regensburg, Germany) and cloned in the expression vector pEXPR103 (iba lifesciences, Goettingen, Germany) in frame with an N-terminal modified mouse Igκ light chain signal peptide. C-terminally, a T4 fibritin foldon trimerization domain (GYIPEAPRDGQAYVRKDGEWVLLSTFL) and a FLAG-tag were added. The original H5 sequence was additionally modified by introducing the aa exchange R345Q in order to prevent cleavage of the full-length H5. To obtain a soluble HA1 subdomain (aa 17 - 345), aa 346 - 530 and the T4 fibritin sequence were deleted from the full-length construct using a restriction-free PCR cloning technique (33). Primer sequences are available upon request.

To produce the H5-nanoluciferase antigen (H5-Nano), the nanoluciferase reporter gene (AFI79290.1) was synthetically produced (GeneArt, Regensburg, Germany) and added c-terminally to the full-length H5 construct. The T4 fibritin sequence was removed and a 12-aa linker sequence (GGSQSDSRGGNG) was inserted upstream of the c-terminal FLAG-tag.

### Protein expression

The recombinant full-length H5 fused to nanoluciferase (H5-Nano), the full-length trimeric H5 and the HA1 subdomain were expressed in Expi293 cells. The cells were grown in Expi293 expression medium (Thermo Fisher Scientific, Waltham, Massachusetts, U.S.) and polycarbonate Erlenmeyer flasks (Corning, New York, U.S.) at 37°C, 8% CO_2_, 125 rpm. Transfection was performed using the ExpiFectamine293 transfection kit (Thermo Fisher Scientific) according to the manufactureŕs instructions and the cell culture supernatant was harvested five days after transfection. The HA1 antigen was purified from the supernatant using ANTI-FLAG® M2 Affinity Gel (A2220; SigmaAldrich/Merck, St. Louis, Missouri, U.S.) as recommended by the manufacturer. The protein was eluted using 100 µg/ml FLAG® peptide (Cat. No. F3290; SigmaAldrich/Merck, St. Louis, Missouri, U.S.). Subsequently, the peptide was removed from the preparation by dialysis against an excess of 1X PBS, pH 7.4. The same procedure was used for the purification of the full-length H5 which was assessed as an alternative capture antigen in the ELISA.

For H5-Nano, the crude supernatant was harvested, aliquoted and stored at −80°C until further use.

### Serum samples

#### Chickens

A total of 476 H5 antibody-negative and -positive serum or plasma samples were obtained during immunization and/or infection studies at the Friedrich-Loeffler-Institut (FLI), Germany. All animal experiments were approved by the animal welfare committee (Landesamt für Landwirtschaft und Fischerei Mecklenburg-Vorpommern (LALLF), Thierfelder Straße 18, 18059 Rostock). Further details are shown in supplementary table 1 and are available upon request.

#### Wild birds and zoo birds

Samples were collected in the course of an outbreak of H5N1 in a zoo in Karlsruhe, Germany in 2022 (34) or were taken from routine diagnostic submissions to the German National Reference Laboratory (NRL) for Avian Influenza (AI) at the FLI (n = 380). An additional 57 samples from wild birds (waterfowl and non-waterfowl) were provided by the Erasmus Medical Centre (EMC), Rotterdam, The Netherlands, from ongoing wild bird surveillance studies.

#### Geese

185 positive and negative sera and plasma samples from geese were obtained during a vaccination study performed at the FLI (35).

#### Cattle

Positive bovine samples were taken from animal trials performed at the FLI (36). A pool of these samples was used as a positive control for the ELISA. Samples that tested negative for anti-influenza A virus antibodies (n = 383) were collected during a previous monitoring study and provided by the German NRL for Bovine Viral Diarrhea (37). A pool of several of these samples served as a negative control for the Nanoluc-ELISA.

#### Swine

Anti-H5 antibody-negative serum samples from domestic pigs (n = 263) and wild boars (n = 48) were collected in Germany during 2009 - 2010 in the frame of previous studies (38, 39). Antibody-positive samples from wild boars (n = 6) were taken from wildlife disease monitoring studies 2024/2025 in Germany (unpublished).

#### Ferret

Anti-H5 antibody-positive samples (n = 6) were obtained during infection studies using A/goose/Germany-BB/ 2020AI00018 (H5N8) at the FLI. Samples from animals infected with H2N2 (n = 4) were provided by the EMC, Rotterdam, The Netherlands (40). Antibody-negative samples (n = 64) were submitted to the NRL for SARS-CoV-2 infections in animals at the FLI for serological investigations in the frame of routine health monitoring in animal facilities.

#### Sheep

714 serum samples from routine diagnostic submissions were provided by the State Office for Agriculture, Food Safety and Fisheries Mecklenburg-Western Pomerania.

#### Wild carnivores

Blood samples (n = 628) were collected 2024 and 2025 in the course of a wildlife disease monitoring project in Mecklenburg-Western Pomerania, Germany (41).

#### Human serum samples

8 anti-H5 antibody-positive sera, collected from individuals previously vaccinated with an adjuvanted inactivated split virion vaccine based on A/Indonesia/5/2005 (clade 2.1) and some of whom had also been vaccinated with a similar vaccine based on A/Vietnam/1194/2004 (clade 1) vaccine two years earlier, were provided by the EMC, Rotterdam, The Netherlands. Sera were collected with informed consent. The Institutional research Review Board of Erasmus MC Rotterdam confirmed that the rules laid down in the Medical Research Involving Human Subjects Act (also known by its Dutch abbreviation WMO) do not apply to the reuse of these samples from the biobank for development of diagnostic methods. Samples from H5-naïve individuals were provided by the Faculty of Medicine and University Hospital Cologne. The sample collection was approved by the Ethics committee of the University of Cologne (EIKIM study protocol, #21-1468) (42).

Further details on all the samples used in the study are provided in supplementary table 1.

### Commercial ELISAs

The ID Screen® Influenza A Antibody Competition Multi-species ELISA (FLUACA-5P; Innovative Diagnostics, Grabels, France), subsequently termed NP-ELISA, was performed according to the instructions of the manufacturer. Cut-off values were applied as recommended: inhibition ≤45%, positive; 45-50%, questionable; ≥50%, negative.

The ID Screen® Influenza H5 Antibody Competition 3.0 Multi-species ELISA (FLUACH5V3-5P; Innovative Diagnostics, Grabels, France), subsequently termed ID Screen H5 ELISA, was performed according to the instructions of the manufacturer. Cut-off values were applied as recommended: inhibition ≤50%, positive; 50-60%, questionable; ≥60%, negative. Some samples from birds and carnivores were tested in the previous version of this ELISA (ID Screen® Influenza A Antibody Competition Multi-species ELISA).

### H5-Nanoluciferase (H5-Nanoluc) ELISA

The H5-Nano detector antigen was expressed in Expi293 cells as described above. Cell culture supernatant was harvested 5 days after transfection and stored at −80°C without further processing. After first thawing, the aliquots were stored at 4°C for up to 4 weeks.

The luciferase activity of the detector antigen per µl of supernatant was quantified using the Nano-Glo® luciferase assay system, (#N1110, Promega, Madison, Wisconsin, U.S.). For this purpose, ten-fold serial dilutions of the supernatant were prepared, mixed with the substrate solution and the bioluminescence was measured in luciferase light units (LU).

High binding white flat bottom ELISA plates (#3922, Corning, New York, U.S.) were coated with 0.2 µg of the HA1 capture antigen per well (2 mg/ml) in 0.1 M carbonate buffer pH 9.6 overnight at 4°C. One well on each plate was left uncoated to control the luciferase activity of the applied H5-Nano dilution. Subsequently, the plates were washed 3 times with washing buffer (1X PBS, 0.1% Tween, pH 7.4) and treated with Liquid Plate Sealer® animal-free (#163 500, CANDOR Bioscience GmbH, Wangen im Allgäu, Germany) as recommended. After that, the plates were sealed and stored desiccated at 4°C until further use.

For the ELISA, serum samples were diluted 1:10 in dilution buffer (1X PBS, 1% Tween, pH 7.4) in a non-binding microtiter plate (#650101, Greiner Bio-One GmbH, Frickenhausen, Germany). The cell culture supernatant containing the H5-Nano antigen was diluted in dilution buffer to 4x 10^6 LU/ml and 50 µl of this dilution was added to each sample (2x 10^5 LU per sample). The mixture of serum and detector antigen was then incubated for 1h at room temperature (RT). In the same time, the pre-coated ELISA plate was blocked with 5% skim milk in 1X PBS for 1h at 37°C. Subsequently, the ELISA plate was washed 3 times with washing buffer and the pre-incubated sample mixture was added for 1h at RT. After washing the plate 6 times with washing buffer, 40 µl of the Nano-Glo substrate solution was added to each well and the luciferase activity (LU) was measured for 2s per well using a Tecan Infinite® F200 PRO plate reader (supplementary figure 1A).

### Modifications for samples from wild terrestrial carnivores

For samples collected post-mortem from wild terrestrial carnivores (red foxes, raccoons, raccoon dogs, European badgers and European pine martens) the original protocol of the Nanoluc ELISA was modified in order to minimize non-specific background signals. The blood samples were mixed 1:10 in dilution buffer and incubated directly on the blocked ELISA plate for 1h at 37°C followed by 5 washes with washing buffer. Subsequently, the H5-Nanoluc detector antigen was diluted in dilution buffer to a concentration of 8 – 10x 10^6 LU/ml and 50 µl was added to each well for 1h at RT. The plate was then washed 6 times with washing buffer and finally, the luciferase activity was measured as described above (supplementary figure 1B).

### Serum neutralization assay (SNT)

The SNT was performed according to a published protocol (43). Briefly, the test samples were heat inactivated at 56°C for 1 hour. Two-fold dilution series of the samples were prepared in cell growth medium supplemented with 2 µg/ml trypsin on 96-well microplates. The test virus was diluted in the same medium to a concentration of 10^3.3 TCID_50_/ml and 50 µl was added to each serum dilution. After incubation for 1h at 37°C the serum/virus mixture was transferred to 96-well plates with confluent MDCK-II cells which had first been washed twice with cell growth medium. The plates were then incubated at 37°C and after 72 hours the read-out was performed by assessing the cytopathic effect. The recombinant test virus carrying the HA gene from A/duck/Vietnam/NCVD-1584/2012 (H5N1, clade 2.3.2.1e) (EPI424984) (44) and the remaining seven gene segments from the A/Puerto Rico/8/1934 (H1N1; PR8) backbone was rescued by reverse genetics according to (45) with slight modifications. Briefly, co-culture of HEK293T and MDCK-II cells was co-transfected with the eight plasmids encoding the desired gene segments and incubated for 72 h. Supernatants were harvested and inoculated into the allantoic cavity of 10-day-old specific-pathogen-free embryonated chicken eggs (Valo BioMedia GmbH, Osterholz-Scharmbeck, Germany). After 72 h, allantoic fluids were harvested, tested for hemagglutination activity and the absence of bacterial contamination. The virus stock was stored at – 80°C until further use.

### Validation methods and statistical analyses

Receiver Operating Characteristic (ROC) analysis was used for determination of cut-off values and assessment of test sensitivity and specificity. Agreement between the commercial ID Screen ELISAs and the H5 Nanoluc assay was calculated using the Cohen’s kappa coefficient (’κ’). Samples that were classified “questionable” by the two commercial ELISAs were excluded from calculation of κ. All statistical analyses were performed using GraphPad Prism 10.0.

## Results

### Protein expression and purification

The H5-HA1 subdomain (aa 17 - 345) used as the capture antigen for the ELISA was expressed in Expi293 cells and FLAG-purified from the supernatant. An optimal coating concentration of 0.2 µg/ well (2 mg/ml) was determined by checkerboard titration using defined antibody-positive and -negative cattle and chicken serum samples.

The H5-Nano detector antigen was expressed at high levels, yielding approximately 1x 10^12 LU/ml luciferase activity in a single 6-well transfection of Expi293 cells which is sufficient for testing more than 1x 10^6 samples. After first thawing, the antigen could be stored at 4°C and for up to two weeks its luciferase activity remained stable. The activity declined by 20 to 30% after one to two months and by 60 to 70% after three to four months of storage at 4°C. In order to maintain reproducibility of the assay, it is therefore crucial to check the luciferase activity of the applied H5-Nano dilution in a separate well of each ELISA plate.

To determine the optimal input concentration of the detector antigen in the ELISA, checkerboard titrations were performed using serially diluted antibody-positive cattle serum samples. It became evident that the highest sensitivity could be achieved by using 10^6 - 10^7 LU of the detector antigen per sample with a limit of detection (LOD) of 1:10240. Using only 10^5 or 10^4 LU showed a comparatively lower sensitivity with LODs of 1:1280 and 1:80, respectively (Figure 1A). In order to achieve an optimal signal-to-noise ratio for the Nanoluc-ELISA, the concentration of the H5-Nano detector was adjusted to 2x 10^5 LU per sample (4x 10^6 LU/ml) or 4-5x 10^5 LU (8-10x 10^6 LU/ml) for the wild carnivore protocol, respectively. Using these concentrations the detection range of the assay for H5-specific antibodies was between 10^2 to 10^6 LU.

**FIG 1:**
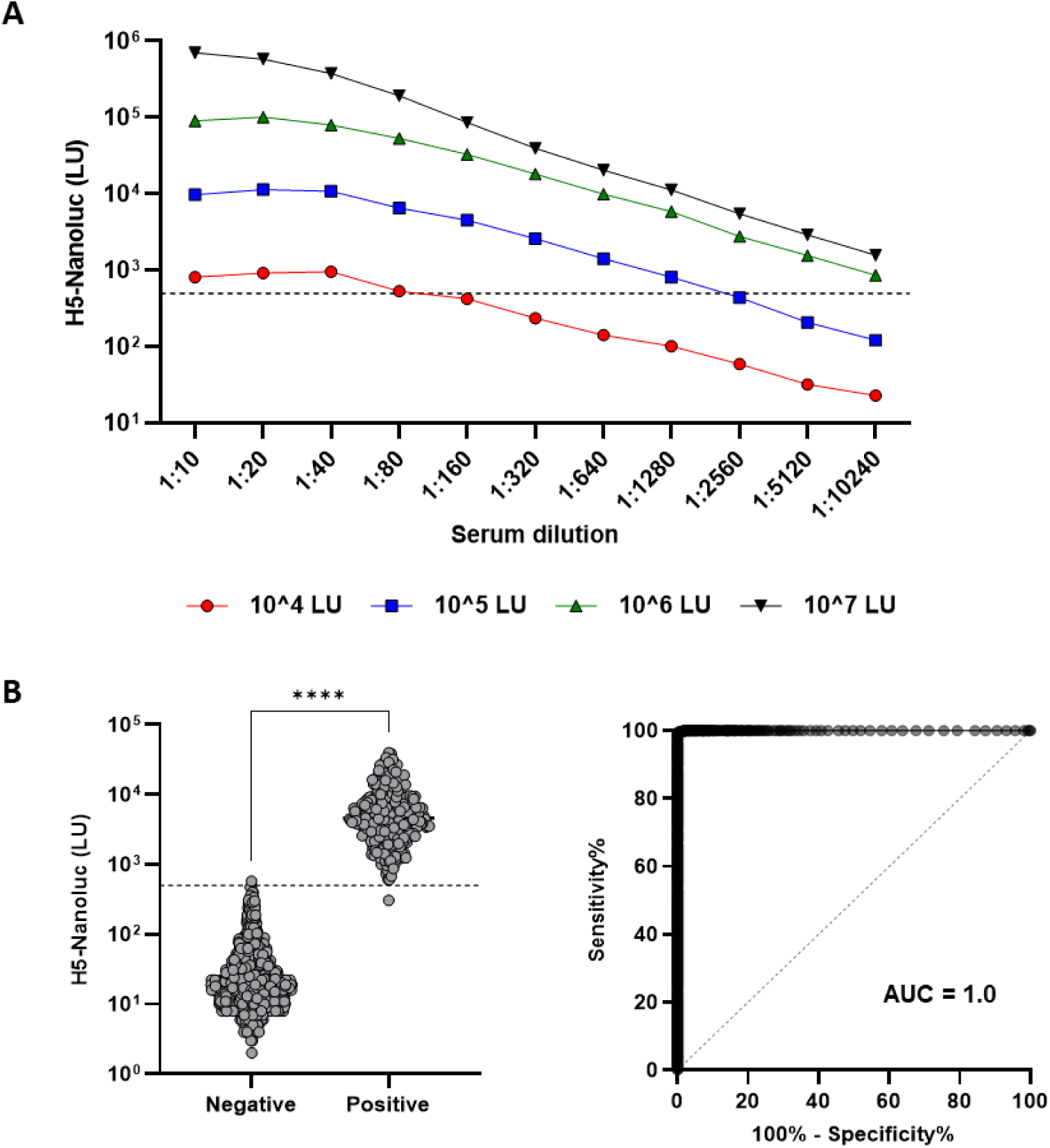
Assay parameters and performance of the H5-Nanoluc assay. (A) Dynamic detection range of the H5-Nanoluc assay dependent on the concentration of the detector antigen. Different dilutions of the H5-Nano antigen (10^4, 10^5, 10^6 or 10^7 LU per sample) were comparatively tested with a serially diluted antibody-positive cattle serum sample. (B) Positive (n = 342) and negative samples (n = 1096) from different mammalian and avian species showed a significantly different ELISA reactivity (unpaired t-test, ****p < 0.0001). The cut-off-value of ≥ 500 LU was determined by ROC analysis and is indicated with a dashed line. ROC analysis yielded an area under the curve (AUC) of 1.0 (95% CI 0.99 - 1.00; p < 0.0001).

### Test performance

A panel of positive (n = 342) and negative (n = 1096) samples from cattle, pigs, wild boars, ferrets, chickens and different bird species (waterfowl and non-waterfowl) was used to choose an optimal cut-off value by ROC analysis. The area under the curve (AUC) was determined 1.0 (95% confidence interval (CI) 0.99 - 1.00; p value < 0.0001). For the selected cut-off value of ≥ 500 LU, the sensitivity was estimated to 99.71% (CI 98.36 - 99.99%) and the specificity to 99.82% (CI 99.34 - 99.97%). Positive sera were defined positive using the commercial ID Screen H5 ELISA as a reference test. All negative samples were tested negative either in the commercial ID Screen NP and/or H5 ELISA (Figure 1B). To assess the within run variation of the ELISA, one sample was tested 40 - 60 times on one plate. In three independent runs, the coefficient of variation (CV%) of the sample was between 6 to 18%. In order to determine the between run variation, 15 samples were tested on 10 different plates on 10 different days. The CV% ranged from 9 to 29% for the individual samples, with higher variations for weakly and very strongly positive samples.

We further addressed the cost-effectiveness of the assay with regard to its applicability for large scale testing. Based on the required materials and the necessary infrastructure, including protein expression and purification, the costs were estimated to approximately 60 € for one plate of 88 samples.

### Detection of H5-specific antibodies in different bird species

H5 clade 2.3.4.4b-antibody-positive samples obtained from vaccinated (n = 102) or infected (n = 6) chickens were used for the initial evaluation of the H5-Nano ELISA and the test allowed a clear and significant (p < 0.0001) distinction of positive and negative samples with a broad dynamic detection range (Figure 2A). Additionally, antibodies were detected in chickens after infection with H5N1 of clade 2.2 or subclade 2.2.2. However, in chickens that were vaccinated with a recombinant herpesvirus of turkey (HVT)-vectored vaccine expressing an H5 subtype of clade 2.2., the H5-Nano assay reacted only with 4 out of 21 samples which were determined H5 antibody-positive by the ID Screen ELISA. A clearly inferior performance was also observed for the detection of antibodies against H5 of clade 1. In chickens vaccinated with a live Newcastle Disease virus (NDV)-vectored vaccine expressing an H5 of clade 1, the ELISA detected 18 out of 41 positive animals and only after homologous challenge infection with H5N1 (A/ck/Viet/P41/05) > 90% of the samples were correctly recognized. Not surprisingly, antibodies of animals infected with a low pathogenicity H5N2 strain (A/duck/Potsdam/1402-6/1986 (H5N2)) not belonging to the A/goose/Guangdong/1/96 lineage were detected only in one out of five animals. Furthermore, no cross-reactivity was observed with antibodies against the HA of subtypes H1, H2 and H6 in samples from experimentally infected chickens. Even though, subtypes H1, H2 and H6 are phylogenetically more closely related to H5 than other subtypes, the measured LU values did not significantly differ from the ones obtained for H5-negative samples (Figure 2A).

**FIG 2:**
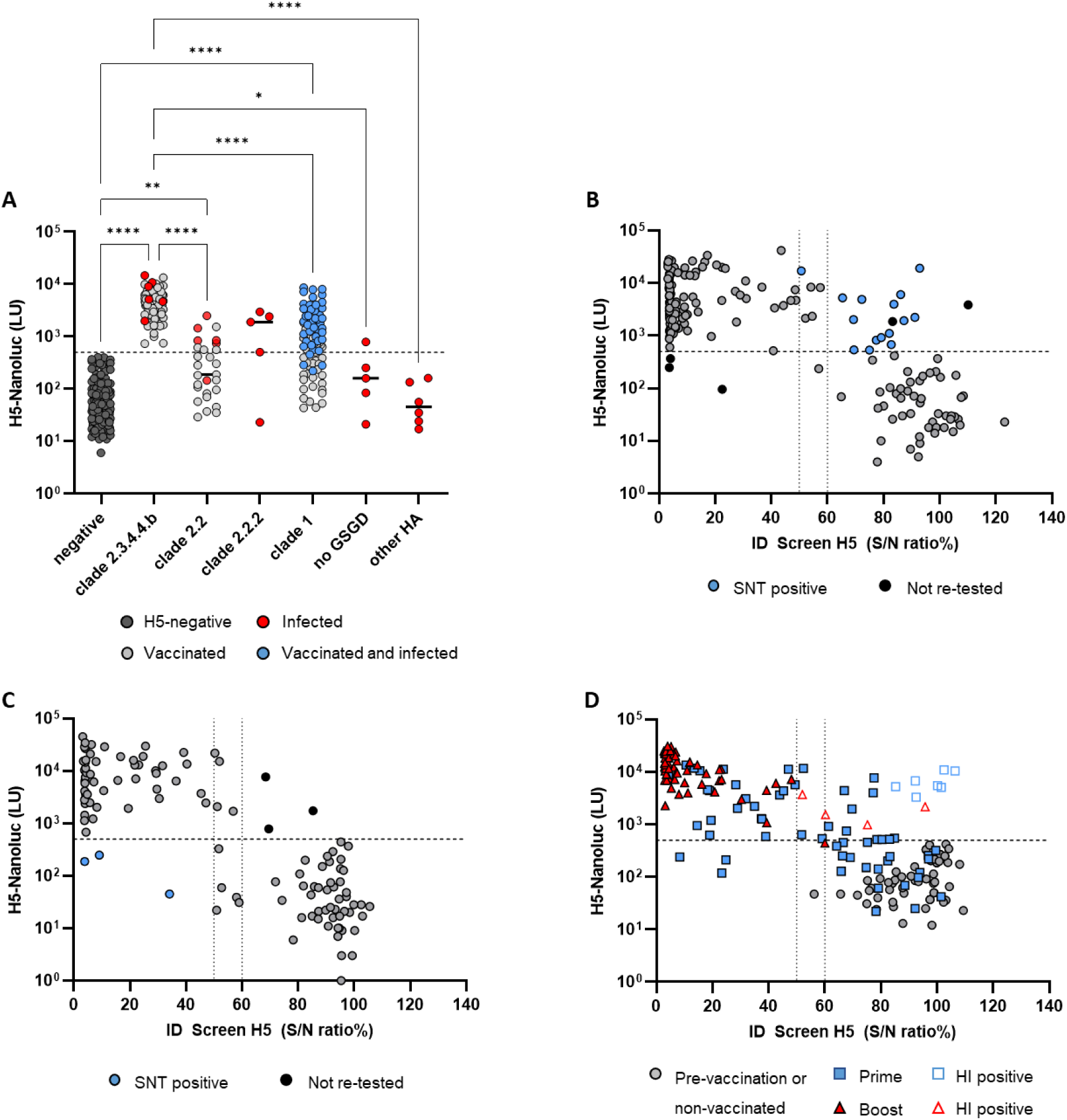
Performance of the Nanoluc-ELISA for samples from avian species. (A) Samples from chickens vaccinated and/or infected with the indicated H5 clades/subclades. Vaccinated animals are shown in grey, infected animals in red. Animals that were vaccinated and subsequently infected are highlighted in blue. “No GSGD” indicates infection with an H5 strain not belonging to the A/goose/Guangdong/1/96 lineage. “Other HA” included antibodies against subtypes H1, H2 and H6. All negative samples were tested negative for anti-influenza A virus antibodies. Differences between groups were analyzed by Kruskal-Wallis test followed by Dunńs multiple comparisons test. Significant differences (p < 0.05) between individual groups vs the negative group and vs the clade 2.3.4.4b group are shown on the graph (****p < 0,0001, **p = 0.0032, *p = 0.019). Samples from waterfowl (B) and from non-waterfowl (C) bird species were comparatively tested with the two H5-specific ELISA formats. Results of the ID Screen H5 ELISA (in S/N% ratio) are plotted on the x-axis and the cut-off values are indicated with dotted lines. Results of the Nanoluc assay (in LU) are plotted on the y-axis, the cut-off value is indicated with a dashed line. Samples colored in blue were tested positive by SNT, samples colored in black could not be re-tested due to lack of material. (D) Comparative analysis of serum samples from geese obtained from non-vaccinated animals and from animals before and after prime and boost immunization with an H5 clade 2.3.4.4b-specific vaccine. Empty symbols depict samples that were previously tested positive in a HI assay (35).

In addition to galliform poultry, the performance of the Nanoluc ELISA was investigated for several other wild and zoo bird species. 193 samples were obtained from wild or captive waterfowl including different duck, goose and swan species (Figure 2B). H5-specific antibodies were detected in 123 and in 120 samples by the ID Screen and the Nanoluc H5 ELISA, respectively. Interestingly, among the 65 samples showing reproducibly negative results in the commercial ELISA, several samples reacted positive in the Nanoluc assay (n = 17; 11 swans, 2 ducks, 4 geese). Based on this outcome, a substantial agreement of the two ELISA formats (Coheńs kappa coefficient k = 0.752, 95% CI 0.651 - 0.853) was determined. However, when samples with inconsistent results were additionally tested in a serum neutralization test (SNT), H5-neutralizing antibodies were detected in the samples yielding positive results in the Nanoluc ELISA which suggests a superior sensitivity compared to the commercial test.

In addition to waterfowl, 122 samples from other wild bird species (gulls, terns, waders, raptors) and captive zoo birds (e.g. peafowl, flamingo, pelicans, cranes, marabou, toucan, ostriches, penguins, owls, macaw, cockatoo and pigeons) were investigated (Figure 2C). The ID Screen H5 and the Nanoluc ELISA detected anti-H5 antibodies in 59 and 56 samples, respectively. Overall, discordant results were observed only for 7 samples (pelicans or gulls) which indicated an almost perfect agreement (k = 0.976, 95% CI: 0.787- 0.965) between the two assays for samples from non-waterfowl bird species.

The suitability of the ELISA to detect antibodies induced upon vaccination, was investigated using samples that were obtained in the frame of two immunization studies in geese (35) (Figure 2D). Within this panel, the agreement of the Nanoluc assay with the commercial ID Screen H5 ELISA was found to be substantial (k = 0.650, 95% CI: 0.522 to 0.777). Notably, disagreements were mainly observed for sera collected after prime but before booster immunization showing positive results in the Nanoluc ELISA whereas the ID Screen ELISA still scored negative (n = 17). Results of a previously performed HI assay confirmed the H5-antibody-positive status of some of these samples (35) (Figure 2D).

### Detection of H5-specific antibodies in mammalian species and humans

In sera of cattle infected with H5N1 clade 2.3.4.4b (36), the Nanoluc ELISA detected H5-specific antibodies starting from 9 days after infection with a significant distinction (p < 0.0001) from negative samples (Figure 3A). Furthermore, the assay demonstrated a superior analytical sensitivity since the limit of detection (LOD) for three different sera was markedly below that of the ID Screen H5 ELISA. Specifically, the LOD of the H5-Nanoluc assay vs the commercial test was found to be 1:80 vs 1:40 for cattle #1, 1:160 vs 1:10 for cattle #2 and 1:640 vs 1:80 for cattle #3 (Figure 3B).

**FIG 3:**
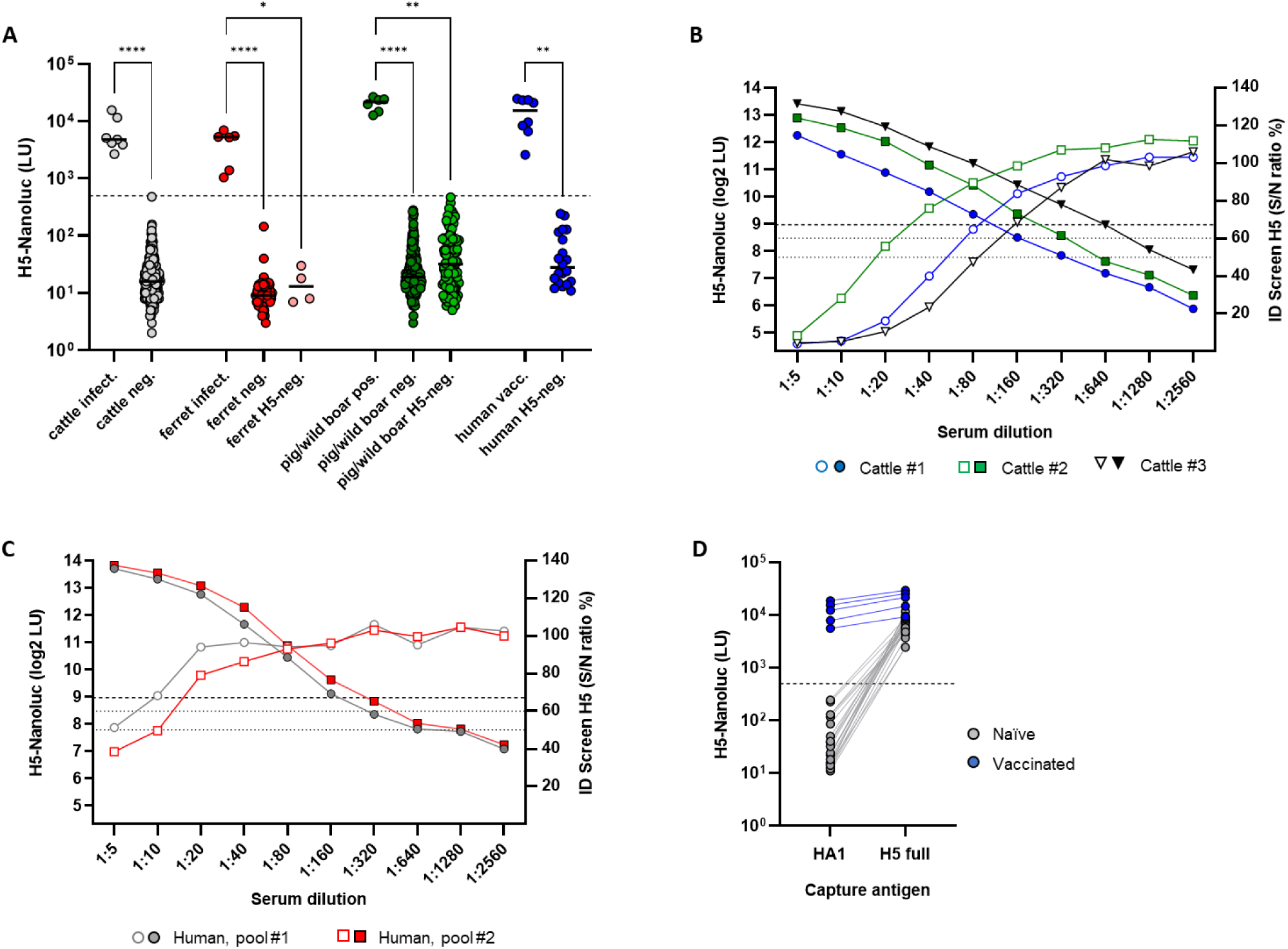
Performance of the Nanoluc ELISA for samples from mammalian species. (A) Samples from cattle, ferrets, pigs, wild boars and humans analyzed with the H5-Nanoluc ELISA. The cut-off value is indicated with a dashed line. Negative samples were tested negative for anti-influenza A virus antibodies, “H5-neg.” samples were tested negative for anti-H5 antibodies but contain antibodies against other HA subtypes (H1, H2 or H3). Statistical differences between antibody-positive and -negative groups of each species were calculated using the Kruskal-Wallis test followed by Dunńs multiple comparisons test and significant differences (p < 0.05) are shown on the graph (****p < 0.0001, **p < 0.01, *p = 0.0135). (B) and (C) Analytical sensitivity and limit of detection of the H5-Nanoluc ELISA in comparison to the ID Screen H5 ELISA for three sera obtained from H5N1-infected cattle (B) or two sample pools from individuals vaccinated with a H5N1-specific vaccine (C). Results of the H5-Nanoluc assay are plotted with bold symbols on the left y-axis, the cut-off value is indicated with a dashed line. Note that LU values are shown as log2 LU in order to allow a distinctive separation of the lines on the graphs. Results of the ID Screen ELISA (in S/N ratio %) are shown with empty symbols on the right y-axis, the cut-off values are indicated with dotted lines. (D) Differentiation of H5 head-specific from cross-reacting anti-stalk antibodies in human serum samples using either the HA1 subdomain or the full-length H5 as a capture antigen in the ELISA. H5-naïve individuals are shown in grey, individuals vaccinated with an H5N1-specific vaccine in blue.

Since no background or non-specific signals were observed during testing of influenza A virus antibody negative cattle sera, additional 714 sera from sheep were tested to probe the test specificity. Among these, 14 NP-antibody-negative samples yielded positive results in the H5 Nanoluc ELISA. However, after re-testing of these samples by SNT, 6 out of 14 showed low levels of neutralizing activity. Excluding these samples from the analysis, a relative test specificity of 98.9 % was determined, which is close to the 99.82% predicted by ROC analysis.

Among samples from domestic pigs and wild boars, high levels of H5-specific antibodies were detected in 6 wild boar samples collected in 2024 in Germany whereas no antibody-positive animals were found in the panels obtained between 2009 and 2010. Interestingly, in about 40% (104 / 263) of the sera originating from domestic pigs, antibodies against influenza A virus were found by the NP-ELISA. However, none of these sera reacted positive in the Nanoluc-assay which underscores its high specificity for H5-specific antibodies (Figure 3A). This finding was confirmed by testing samples from experimentally infected ferrets. Antibodies raised upon infection with H5N1 reacted positively, showing high LU signals whereas no signals were observed for naïve ferrets or ferrets infected with H2N2 viruses (Figure 3A).

Human samples collected from individuals immunized with an H5N1-specific vaccine reacted highly positive in the Nanoluc assay. Testing dilution series of two antibody-positive serum pools determined a LOD of 1:160 for the Nanoluc ELISA compared to an LOD of 1:10 in the commercial ELISA, which demonstrates a superior analytical sensitivity of the novel assay (Figure 3C). All serum samples from H5N1-naïve individuals were tested negative, despite containing H1- and H3-specific antibodies (42). In addition, the previously identified cross-reactive antibodies targeting the HA stalk domain (42) were not detected using the HA1 capture antigen whereas their presence could be confirmed by using the full-length H5 as the capture antigen in the ELISA (Figure 3D). This offers the possibility to differentiate cross-reacting stalk antibodies from anti-H5 head specific antibodies by using both capture antigens in parallel.

For testing samples obtained post-mortem from wild carnivores it was necessary to modify the original ELISA protocol due to high non-specific background signals. This problem was overcome by incubating the diluted samples on the coated plate first, followed by a washing step and subsequent adding of the H5-Nano detector antigen instead of performing the original liquid-phase co-incubation. Positive (n = 127) and negative (n = 443) samples from red foxes, raccoons, raccoon dogs, European badgers and European pine martens (41) were used to evaluate the diagnostic performance of this adapted protocol. ROC analysis yielded an AUC of 0.9926 (95% CI: 0.9867 - 0.9984; p value < 0.0001). Applying a cut-off value of ≥ 380 LU the test sensitivity was estimated to 95% (CI 90.08 – 97.82%) and the specificity to 98% (CI 96.18 – 98.93%) (Figure 4A).

**FIG 4:**
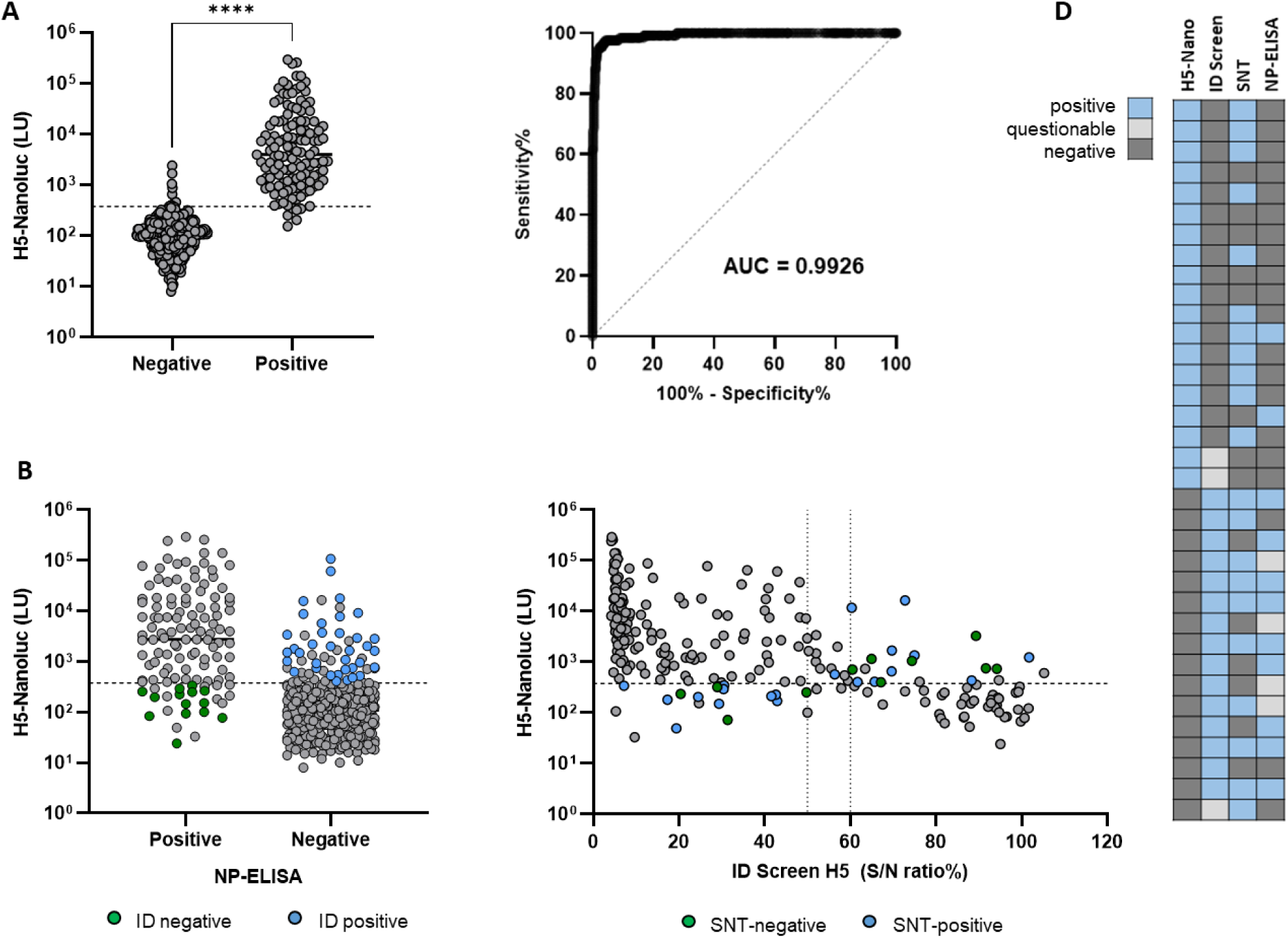
Performance of the Nanoluc ELISA for samples from wild terrestrial carnivores. (A) Positive (n = 127) and negative (n = 443) samples showed a significantly different ELISA reactivity (unpaired t-test, ****p < 0.0001). The cut-off-value of ≥ 380 LU was determined by ROC analysis and is indicated with a dashed line. ROC analysis yielded an area under the curve (AUC) of 0.9926 (95% CI 0.9867 – 0.9984, p < 0.0001). (B) Detection of H5-specific antibodies by the Nanoluc ELISA in samples pre-tested for antibodies against the influenza nucleoprotein using the commercial NP-ELISA. The cut-off value is indicated with a dashed line. Samples highlighted in green or in blue were tested negative or positive, respectively in the ID Screen H5 ELISA. (C) Comparison of the H5-Nanoluc assay and the ID Screen H5 ELISA for samples tested in both assays in parallel. Results of the ID Screen H5 ELISA (in S/N ratio %) are plotted on the x-axis and the cut-off values are indicated with dotted lines. Results of the Nanoluc assay (in LU) are plotted on the y-axis, the cut-off value is indicated with a dashed line. Samples colored in blue were tested positive by SNT, samples in green were negative. (D) Comparison of test results from four independent serological assays for a subset of samples with disagreeing results in the H5-specific diagnostic tests. Each row represents one sample.

Upon pre-screening with the NP-specific ELISA, 121 out of 619 carnivore samples tested positive for influenza A virus specific antibodies. The Nanoluc assay subsequently found H5-specific antibodies in 79% of these samples (96 / 121). It can be assumed, that antibodies in the remaining samples are directed against other H5 clades or HA subtypes since they were mostly also classified negative or questionable by the ID Screen H5 ELISA. Interestingly, several anti-NP-negative samples reacted positive in both, the Nanoluc and the ID Screen ELISA which might indicate a superior sensitivity of these assays for the detection of subtype-specific antibodies (Figure 4B). However, by directly comparing the results of samples tested in both H5-ELISAs in parallel (n = 215) only a moderate agreement was determined (k = 0.532, 95% CI 0.403 - 0.661) and for several samples clear discrepancies became obvious between the two assays (Figure 4C).

In order to investigate this in more detail, we used the SNT as an additional, independent method to analyze these samples. Comparing the results of the three different ELISAs and the SNT revealed inconsistencies among all the assays which implies that the quality of the test material has a considerable impact on the test performance (Figure 4D).

## Discussion

The ongoing panzootic demonstrates an unprecedented spread and rapid evolution of the H5N1 HPAI virus with numerous outbreaks and massive die-offs in wild and domestic animals never reported before. Rising numbers of spillover infections in mammals indicate an elevated risk of zoonotic transmission, emphasizing the need for continued monitoring and targeted preparedness measures. Thus, an increased serological surveillance is critical to identify new host species and to uncover potential paths of silent spread. For this purpose, robust assays with no restriction to selected species are urgently needed.

In the present study, a novel H5-specific ELISA was developed based on the previously described LIPS technology which has proven to be a highly sensitive antibody detection method for diagnosis of a broad variety of autoimmune and infectious diseases (46, 47). A LIPS assay for detection of H5-specific antibodies using HA1 and HA2 detector-antigens has previously been described (48). However, the original LIPS platform is based on the binding of antibody-antigen complexes to protein G beads which is not applicable for all species, most notably from avian origin. In contrary, a double antigen ELISA format allows a species-independent detection of antibodies against influenza A virus as it has been shown before (49). However, the validation of such a broadly applicable test is challenging due to the lack of reference samples and gold standard tests for many species, especially wildlife (50, 51). Among several previously developed ELISAs, variations of sensitivity, specificity and accuracy have been described for different bird species, in particular among waterfowl (18, 23, 52, 53). The H5-Nanoluc ELISA allowed a sensitive, specific and homogeneous detection of H5-specific antibodies in a broad variety of bird species in a good agreement with the ID Screen H5 ELISA. However, for samples from ducks, geese and swans, the Nanoluc assay demonstrated a superior sensitivity compared to the commercial ELISA. One possible explanation for this finding could be the distinct antibody repertoire in non-galliform birds. In addition to the full-length IgY antibody, ducks additionally produce a smaller version of IgY lacking the Fc region (IgYΔFc) which becomes predominant in the serum at later stages of the immune response (54, 55).

Using the double-antigen ELISA format, antibodies are detected by simultaneous binding to their specific antigen, which means that all immunoglobulin isotypes including IgYΔFc fragments can be efficiently captured. In the competitive ID Screen ELISA, the smaller IgY versions might not be able to compete sufficiently with the labeled detector antibody which can lead to false-negative test results. The superior performance of the Nanoluc ELISA for samples from waterfowl was further underscored by testing samples from vaccinated geese. In several animals, the assay detected antibodies already after prime immunization whereas the ID Screen ELISA mostly yielded negative results before the booster vaccination. Its outstanding sensitivity to detect even low levels of H5-specific antibodies makes the Nanoluc format a valuable tool for monitoring vaccination campaigns, especially in ducks and geese. Nevertheless, these findings also implicate that the high sensitivity of the test could lead to a certain amount of false-positive test results. Thus, positive reactions of samples tested negative for anti-influenza A virus antibodies in a NP-specific ELISA should be assessed carefully and might require additional confirmation.

With the emergence of H5N1 2.3.4.4b viruses serological screening and monitoring of mammalian species has become increasingly important due to the frequent spillover events and adaptation to new hosts (1, 2). The Nanoluc ELISA showed a good diagnostic performance for samples from cattle, domestic pigs, wild boars, ferrets and humans. Particularly for samples from cattle and vaccinated humans, the newly developed assay demonstrated a superior analytical sensitivity and lower limits of detection compared to the commercial H5 ELISA. This could be explained by the fact that different species use varying strategies to create their individual antibody repertoires (56). Consequently, some species might develop antibodies preferentially targeting epitopes on H5 which are not covered by the detector antibody in the competition ELISA. In contrast, the liquid-phase Nanoluc ELISA allows binding of antibodies to all accessible epitopes on the antigen, independent of species-specific variations. Accordingly, the test was suitable for the detection of H5-specific antibodies in several wild carnivore species but only in a moderate agreement with the ID Screen ELISA. However, it has to be considered that the respective samples were collected postmortem from carcasses with autolytic processes already started. Thus, discordant test results are likely caused by assay-specific susceptibilities for certain inhibitors which also explains the necessity to modify the protocol of the H5-Nanoluc ELISA. The lack of suitable reference samples and a true gold standard test makes the interpretation of the data and the observed disagreements challenging. Therefore, we additionally analyzed some of the samples by SNT and compared the outcome with the results of the two H5-specific and the NP-specific ELISA. Inconsistencies between all assays became obvious and thus confirmed, that the quality of the sample matrix indeed has an impact on the test performance, independent of the test format. However, these results also show, that using a combination of the two different H5-specific assays can compensate some weaknesses of each test, allowing to reliably determine the prevalence of H5-specific antibodies among populations of wild carnivores. Furthermore, a complementary application would improve the diagnostic sensitivity for the detection of H5 subtype-specific antibodies. In contrast to the broad reactivity of the ID Screen ELISA, the Nanoluc assay revealed a distinct specificity for antibodies against H5 of clade 2.3.4.4b, whereas antibodies against other H5 clades, especially clade 1, could not be reliably detected. This finding coincides with the previous observation that H5 2.3.4.4b-specific antibodies bind strongly to the homotypic antigen but only weakly to H5 proteins of other clades indicating that the antibodies target certain clade-specific epitopes (57).

It also became obvious, that the selection and design of the capture and detector antigen play a crucial role for the specificity of the Nanoluc assay. Using the full-length H5 as a detector antigen allows binding and detection of all antibodies elicited against both the H5 head and stalk domain. However, in combination with HA1 serving as the capture antigen, the detection is limited to antibodies specifically targeting the H5 head domain which requires an exposure to this particular antigen. As a result, the Nanoluc ELISA did not react with antibodies against HA subtypes H2 and H6, whereas this could repeatedly be observed in previously developed assays (19, 29). This high specificity was further underscored by the lack of cross-reactivity with antibodies against H1 and H3 in humans or in samples from pigs and wild boars. Based on these observations, the Nanoluc-ELISA represents a valuable tool for the detection of past H5-infections in pigs. Even though, previous studies demonstrated a low susceptibility of pigs to infections with H5N1, including the bovine-derived H5N1 B3.13 (58, 59), they represent a potential host for the generation of novel virus reassortants with a possible pre-pandemic risk. Thus, an increased surveillance is crucial for early awareness and to be able to prevent transmission (16).

The H5-Nanoluc ELISA could further be helpful for the monitoring of future vaccination strategies with an H5N1-specific vaccine (60). In previous studies it was shown that broadly cross-reacting antibodies directed against the conserved HA stalk domain are produced following vaccination with seasonal influenza vaccines. Consequently, detectable levels of cross-neutralizing antibodies can also be found in individuals that have not been exposed to H5N1 (42, 61, 62). These pre-existing anti-stalk antibodies are not detected by the Nanoluc ELISA since the HA1 subdomain is used as the capture antigen. Therefore, the assay does not only allow to verify the generation of anti-H5 antibodies after H5-vaccination but can be used to identify H5-naïve individuals.

## Conclusions

We could show, that the newly developed, luciferase-based double antigen ELISA allows a highly sensitive and specific detection of H5-specific antibodies in a broad range of avian and mammalian species including humans. Compared to a commercially available H5-specific ELISA, the test showed a superior performance with regard to a species-independent application, especially in waterfowl and in mammals. The assay further proved high specificity for antibodies targeting the H5 clade 2.3.4.4b head domain and no cross-reactivity with other HA subtypes was detected. Therefore, it represents a valuable contribution to existing serological tests and in combination with other assay formats it could markedly improve serological diagnostic tools for the detection of H5-specific antibodies across different host species.

## Funding

Funded by the European Union under grant agreement (101084171) - (Kappa-Flu). Views and opinions expressed are however those of the author(s) only and do not necessarily reflect those of the European Union or REA. Neither the European Union nor the granting authority can be held responsible for them.

## Acknowledgments

We thank Cornelia Illing, Bianka Hillmann, Mareen Grawe, Julien Schäfer, Alrik-Markis Kunisch, Martina Abs and Franziska Rachel for excellent technical assistance. We thank Uwe Nitz, Clemens Landmesser, Ulf Damrau (†), Sophia Ziegler, Jonas Heck, Ralf Redmer, Thomas Pieper and Marco Beerbohm for support with transport and during necropsies. We thank all hunters contributing to this study, especially the hunters organized by the “Jagdverband Rügen-Hiddensee”. We thank Professor Florian Klein and his team at the Institute of Virology, University Hospital Cologne, for providing serum samples from H5N1-naïve individuals for this study.

---

**SUPPL. FIG 1:**
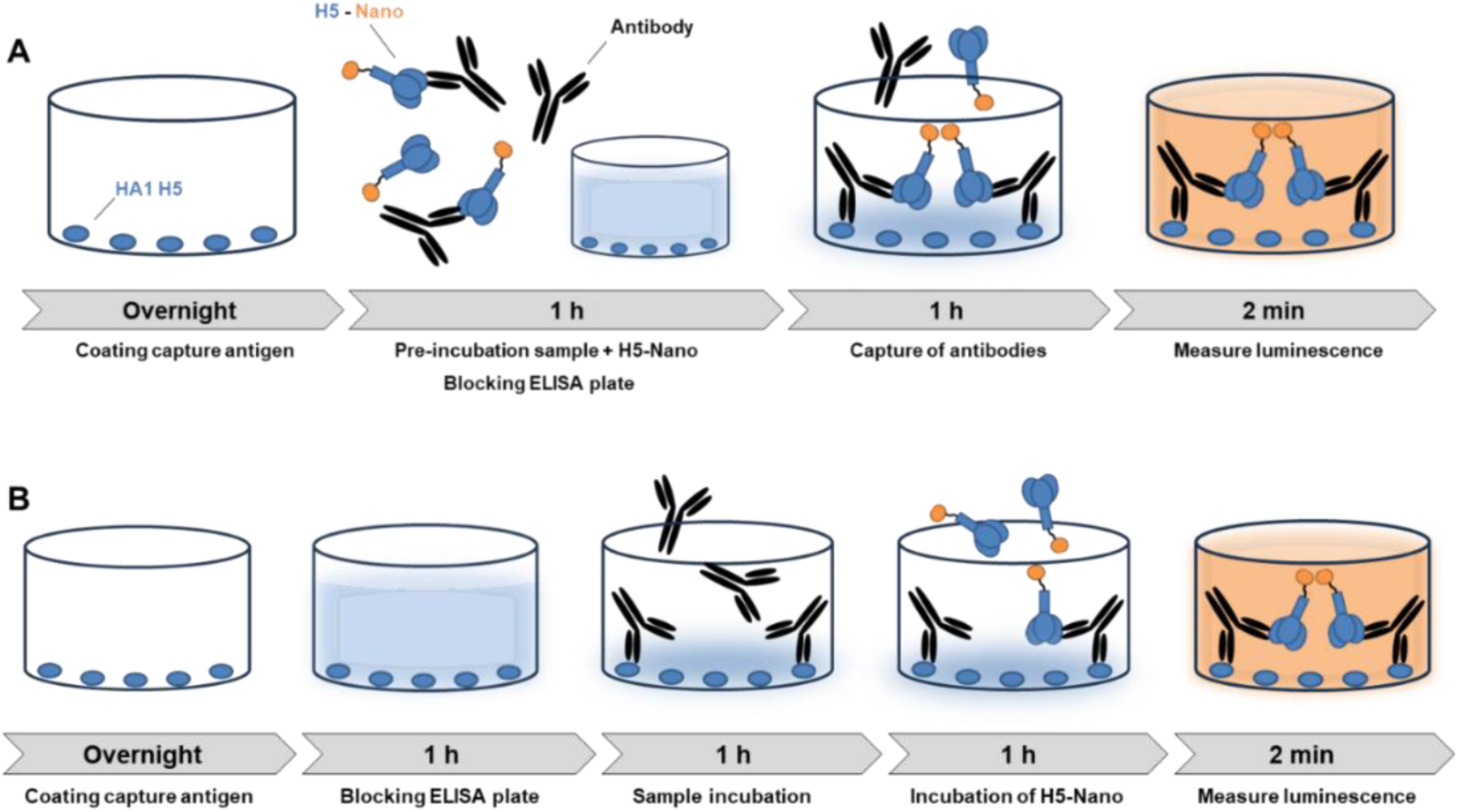
Concept and main steps of the H5 Nanoluc assay. (A) For the original ELISA protocol plates are coated overnight with the HA1 capture antigen. For the assay, samples are diluted 1:10 in dilution buffer and pre-incubated with the H5-Nano detector antigen for 1h at RT. In parallel, the ELISA plate is blocked with 5% skim milk at 37°C. After washing, the sample-detector mixture is added to the ELISA plate and incubated for 1h at RT. After six washing steps, Nano-Glo substrate solution is added to each well and the luciferase activity is recorded. (B) Modified protocol of the Nanoluc assay for samples from wild carnivores with sequential incubation of the diluted test sample and the H5-Nano detector antigen.

**SUPPL. TABLE 1:** Details of samples used for evaluation of the H5-Nanoluc ELISA.

| Species | Number | H5 antibody status | H5 infection / vaccination status | Source of samples |
| --- | --- | --- | --- | --- |
| Chickens | 6 | positive | infected with H5N1 (A/chi/Ger-SH/AI08298/2021), H5 clade 2.3.4.4b | Infection studies performed at the FLI |
|  | 6 | positive | infected with H5N1 (A/chi/Viet/P41/05), clade 2.2 |  |
|  | 5 | positive | infected with H5N1 (A/Cygnus cygnus/Germany/R65/2006), subclade 2.2.2 |  |
|  | 5 | positive | infected with A/duck/Potsdam/1402-6/1986 (H5N2), no GSGD |  |
|  | 102 | positive | vaccinated with a recombinant Newcastle Disease Virus (NDV) expressing H5 clade 2.3.4.4b and H5 clade 1 | Vaccination study performed at the FLI |
|  | 122 | positive / negative | vaccinated with a recombinant NDV expressing an H5 clade 1 and infected with H5N1 (A/chi/Viet/P41/05) | Vaccination and challenge infection study performed at the FLI |
|  | 36 | positive / negative | vaccinated with a recombinant herpesvirus of turkey (HVT)-vectored vaccine expressing an H5 of clade 2.2 | Vaccination study performed at the FLI |
|  | 188 | negative | naïve, non-infected or non-vaccinated | Collected from trials mentioned above |
|  | 6 | negative | infected with H6N2, H2N5, H2N9 or H1Nx | Infections studies performed at the FLI |
| Zoo birds | 316 | positive / negative | infected with H5N1 / naïve | Zoo Karlsruhe, Germany (34) |
| Gulls, swans | 20, 44 | positive / negative | infected with H5N1 / naïve | Routine diagnostic submission to the NRL for AI at the FLI |
| Wild birds | 57 | positive / negative | infected with H5N1 / naïve | Provided by the Erasmus Medical Center (EMC), Rotterdam, The Netherlands from ongoing wild bird surveillance studies. |
| Geese | 185 | positive / negative | vaccinated with an H5N1-specific vaccine, H5 clade 2.3.4.4b | Vaccination study performed at the FLI (35) |
| Ferrets | 6 | positive | infected with H5N1 | Infection study performed at the FLI (unpublished data) |
|  | 4 | negative | infected with H2N2 | Provided by the EMC, Rotterdam, The Netherlands (40) |
|  | 64 | negative | naïve | Routine diagnostic submissions to the NRL for SARS-CoV-2 infections in animals at the FLI |
| Sheep | 714 | negative | naïve | Routine diagnostic submissions to the State Office for Agriculture, Food Safety and Fisheries Mecklenburg-Western Pomerania 2024 and 2025 |
| Cattle | 7 | positive | infected with H5N1 | Infection study performed at the FLI (36) |
|  | 383 | negative | naïve | Surveillance study (37) |
| Domestic pigs | 159 | negative | naïve | Surveillance study (38) |
|  | 104 | negative | previous infection with H1Nx |  |
| Wild boars | 48 | negative | naïve | Surveillance study (39) |
|  | 6 | positive | infected with H5N1 | Wildlife disease monitoring project in Germany 2024/25 (unpublished) |
| Carnivores | 628 | positive / negative | infected with H5N1 / naïve | Wildlife disease monitoring project in Mecklenburg-Western Pomerania, Germany (41) |
| Humans | 8 | positive | vaccinated with H5N1-specific vaccine | Provided by the EMC, Rotterdam, The Netherlands as described in Materials and Methods. |
|  | 20 | negative | naïve | Provided by the Faculty of Medicine and University Hospital Cologne as described in Materials and Methods (42) |

## References

1. Graziosi G, Lupini C, Catelli E, Carnaccini S. 2024. Highly Pathogenic Avian Influenza (HPAI) H5 Clade 2.3.4.4b Virus Infection in Birds and Mammals. Animals (Basel) 14.

2. Peacock TP, Moncla L, Dudas G, VanInsberghe D, Sukhova K, Lloyd-Smith JO, Worobey M, Lowen AC, Nelson MI. 2025. The global H5N1 influenza panzootic in mammals. Nature 637:304–313.

3. Bellido-Martin B, Rijnink WF, Iervolino M, Kuiken T, Richard M, Fouchier RAM. 2026. Evolution, spread and impact of highly pathogenic H5 avian influenza A viruses. Nat Rev Microbiol 24:45–60.

4. FAO. 2025. Global Avian Influenza Viruses with Zoonotic Potential situation update, *on* FAO. https://www.fao.org/animal-health/situation-updates/global-aiv-with-zoonotic-potential/bird-species-affected-by-h5nx-hpai. Accessed 2025.

5. Plaza PI, Gamarra-Toledo V, Eugui JR, Lambertucci SA. 2024. Recent Changes in Patterns of Mammal Infection with Highly Pathogenic Avian Influenza A(H5N1) Virus Worldwide. Emerg Infect Dis 30:444–452.

6. Kuiken T, Vanstreels RET, Banyard A, Begeman L, Breed A, Dewar M, Fijn R, Serafini PP, Uhart M, Wille M. 2025. Emergence, spread, and impact of high-pathogenicity avian influenza H5 in wild birds and mammals of South America and Antarctica. Conserv Biol doi:10.1111/cobi.70052:e70052.

7. Leguia M, Garcia-Glaessner A, Munoz-Saavedra B, Juarez D, Barrera P, Calvo-Mac C, Jara J, Silva W, Ploog K, Amaro L, Colchao-Claux P, Johnson CK, Uhart MM, Nelson MI, Lescano J. 2023. Highly pathogenic avian influenza A (H5N1) in marine mammals and seabirds in Peru. Nat Commun 14:5489.

8. Uhart MM, Vanstreels RET, Nelson MI, Olivera V, Campagna J, Zavattieri V, Lemey P, Campagna C, Falabella V, Rimondi A. 2024. Epidemiological data of an influenza A/H5N1 outbreak in elephant seals in Argentina indicates mammal-to-mammal transmission. Nat Commun 15:9516.

9. Tomas G, Marandino A, Panzera Y, Rodriguez S, Wallau GL, Dezordi FZ, Perez R, Bassetti L, Negro R, Williman J, Uriarte V, Grazioli F, Leizagoyen C, Riveron S, Coronel J, Bello S, Paez E, Lima M, Mendez V, Perez R. 2024. Highly pathogenic avian influenza H5N1 virus infections in pinnipeds and seabirds in Uruguay: Implications for bird-mammal transmission in South America. Virus Evol 10:veae031.

10. Pardo-Roa C, Nelson MI, Ariyama N, Aguayo C, Almonacid LI, Gonzalez-Reiche AS, Munoz G, Ulloa M, Avila C, Navarro C, Reyes R, Castillo-Torres PN, Mathieu C, Vergara R, Gonzalez A, Gonzalez CG, Araya H, Castillo A, Torres JC, Covarrubias P, Bustos P, van Bakel H, Fernandez J, Fasce RA, Johow M, Neira V, Medina RA. 2025. Cross-species and mammal-to-mammal transmission of clade 2.3.4.4b highly pathogenic avian influenza A/H5N1 with PB2 adaptations. Nat Commun 16:2232.

11. Aguero M, Monne I, Sanchez A, Zecchin B, Fusaro A, Ruano MJ, Del Valle Arrojo M, Fernandez-Antonio R, Souto AM, Tordable P, Canas J, Bonfante F, Giussani E, Terregino C, Orejas JJ. 2023. Highly pathogenic avian influenza A(H5N1) virus infection in farmed minks, Spain, October 2022. Euro Surveill 28.

12. Kareinen L, Tammiranta N, Kauppinen A, Zecchin B, Pastori A, Monne I, Terregino C, Giussani E, Kaarto R, Karkamo V, Lahteinen T, Lounela H, Kantala T, Laamanen I, Nokireki T, London L, Helve O, Kaariainen S, Ikonen N, Jalava J, Kalin-Manttari L, Katz A, Savolainen-Kopra C, Lindh E, Sironen T, Korhonen EM, Aaltonen K, Galiano M, Fusaro A, Gadd T. 2024. Highly pathogenic avian influenza A(H5N1) virus infections on fur farms connected to mass mortalities of black-headed gulls, Finland, July to October 2023. Euro Surveill 29.

13. Caserta LC, Frye EA, Butt SL, Laverack M, Nooruzzaman M, Covaleda LM, Thompson AC, Koscielny MP, Cronk B, Johnson A, Kleinhenz K, Edwards EE, Gomez G, Hitchener G, Martins M, Kapczynski DR, Suarez DL, Alexander Morris ER, Hensley T, Beeby JS, Lejeune M, Swinford AK, Elvinger F, Dimitrov KM, Diel DG. 2024. Spillover of highly pathogenic avian influenza H5N1 virus to dairy cattle. Nature 634:669–676.

14. Mostafa A, Naguib MM, Nogales A, Barre RS, Stewart JP, Garcia-Sastre A, Martinez-Sobrido L. 2024. Avian influenza A (H5N1) virus in dairy cattle: origin, evolution, and cross-species transmission. mBio 15:e0254224.

15. Burrough ER, Magstadt DR, Petersen B, Timmermans SJ, Gauger PC, Zhang J, Siepker C, Mainenti M, Li G, Thompson AC, Gorden PJ, Plummer PJ, Main R. 2024. Highly Pathogenic Avian Influenza A(H5N1) Clade 2.3.4.4b Virus Infection in Domestic Dairy Cattle and Cats, United States, 2024. Emerg Infect Dis 30:1335–1343.

16. Krammer F, Hermann E, Rasmussen AL. 2025. Highly pathogenic avian influenza H5N1: history, current situation, and outlook. J Virol 99:e0220924.

17. Niu Q, Jiang Z, Wang L, Ji X, Baele G, Qin Y, Lin L, Lai A, Chen Y, Veit M, Su S. 2025. Prevention and control of avian influenza virus: Recent advances in diagnostic technologies and surveillance strategies. Nat Commun 16:3558.

18. Dlugolenski D, Hauck R, Hogan RJ, Michel F, Mundt E. 2010. Production of H5-specific monoclonal antibodies and the development of a competitive enzyme-linked immunosorbent assay for detection of H5 antibodies in multiple species. Avian Dis 54:644–9.

19. Postel A, Ziller M, Rudolf M, Letzel T, Ehricht R, Pourquier P, Dauber M, Grund C, Beer M, Harder TC. 2011. Broad spectrum reactivity versus subtype specificity-trade-offs in serodiagnosis of influenza A virus infections by competitive ELISA. J Virol Methods 173:49–59.

20. Moreno A, Lelli D, Brocchi E, Sozzi E, Vinco LJ, Grilli G, Cordioli P. 2013. Monoclonal antibody-based ELISA for detection of antibodies against H5 avian influenza viruses. J Virol Methods 187:424–30.

21. Saczynska V, Florys-Jankowska K, Porebska A, Cecuda-Adamczewska V. 2021. A novel epitope-blocking ELISA for specific and sensitive detection of antibodies against H5-subtype influenza virus hemagglutinin. Virol J 18:91.

22. Prabakaran M, Ho HT, Prabhu N, Velumani S, Szyporta M, He F, Chan KP, Chen LM, Matsuoka Y, Donis RO, Kwang J. 2009. Development of epitope-blocking ELISA for universal detection of antibodies to human H5N1 influenza viruses. PLoS One 4:e4566.

23. Sullivan HJ, Blitvich BJ, VanDalen K, Bentler KT, Franklin AB, Root JJ. 2009. Evaluation of an epitope-blocking enzyme-linked immunosorbent assay for the detection of antibodies to influenza A virus in domestic and wild avian and mammalian species. J Virol Methods 161:141–6.

24. Hochman O, Xu W, Yang M, Yang C, Ambagala A, Rogiewicz A, Wang JJ, Berhane Y. 2023. Development and Validation of Competitive ELISA for Detection of H5 Hemagglutinin Antibodies. Poultry 2:349–362.

25. Zhao S, Schuurman N, Tieke M, Quist B, Zwinkels S, van Kuppeveld FJM, de Haan CAM, Egberink H. 2020. Serological Screening of Influenza A Virus Antibodies in Cats and Dogs Indicates Frequent Infection with Different Subtypes. J Clin Microbiol 58.

26. Li ZN, Trost JF, Weber KM, LeMasters EH, Nasreen S, Esfandiari J, Gunasekera AH, McCausland M, Sturm-Ramirez K, Wrammert J, Gregory S, Veguilla V, Stevens J, Miller JD, Katz JM, Levine MZ. 2017. Novel multiplex assay platforms to detect influenza A hemagglutinin subtype-specific antibody responses for high-throughput and in-field applications. Influenza Other Respir Viruses 11:289–297.

27. Cordero-Ortiz M, Magtoto R, Cauwels B, Baum DH, Arruda B, Gorden PJ, Magstadt DR, Hernandez J, Gimenez-Lirola LG. 2025. Fluorescent Microsphere Immunoassay for Isotype-Specific H5N1 Antibody Detection in Serum and Milk Samples From Dairy Cattle: A Tool for Epidemiological Surveillance. J Med Virol 97:e70321.

28. Chestakova IV, van der Linden A, Bellido Martin B, Caliendo V, Vuong O, Thewessen S, Hartung T, Bestebroer T, Dekker J, Jonge Poerink B, Grone A, Koopmans M, Fouchier R, van den Brand JMA, Sikkema RS. 2023. High number of HPAI H5 virus infections and antibodies in wild carnivores in the Netherlands, 2020-2022. Emerg Microbes Infect 12:2270068.

29. Freidl GS, de Bruin E, van Beek J, Reimerink J, de Wit S, Koch G, Vervelde L, van den Ham HJ, Koopmans MP. 2014. Getting more out of less--a quantitative serological screening tool for simultaneous detection of multiple influenza A hemagglutinin-types in chickens. PLoS One 9:e108043.

30. Burbelo PD, Goldman R, Mattson TL. 2005. A simplified immunoprecipitation method for quantitatively measuring antibody responses in clinical sera samples by using mammalian-produced Renilla luciferase-antigen fusion proteins. BMC Biotechnol 5:22.

31. Burbelo PD, Ching KH, Klimavicz CM, Iadarola MJ. 2009. Antibody profiling by Luciferase Immunoprecipitation Systems (LIPS). J Vis Exp doi:10.3791/1549.

32. Hall MP, Unch J, Binkowski BF, Valley MP, Butler BL, Wood MG, Otto P, Zimmerman K, Vidugiris G, Machleidt T, Robers MB, Benink HA, Eggers CT, Slater MR, Meisenheimer PL, Klaubert DH, Fan F, Encell LP, Wood KV. 2012. Engineered luciferase reporter from a deep sea shrimp utilizing a novel imidazopyrazinone substrate. ACS Chem Biol 7:1848–57.

33. Geiser M, Cebe R, Drewello D, Schmitz R. 2001. Integration of PCR fragments at any specific site within cloning vectors without the use of restriction enzymes and DNA ligase. Biotechniques 31:88-+.

34. Anne Günther ea. 2025. Segregate, Test, Observe and Persevere’ (STOP): Strengthening ex situ breeding programmes for biodiversity in zoos amid highly pathogenic avian influenza threats – a case approach, under revision.

35. Piesche R, Cazaban C, Frizzo da Silva L, Ramírez-Martínez L, Hufen H, Beer M, Harder T, Grund C. 2025. Immunogenicity and Protective Efficacy of Five Vaccines Against Highly Pathogenic Avian Influenza Virus H5N1, Clade 2.3.4.4b, in Fattening Geese. Vaccines 13:399.

36. Halwe NJ, Cool K, Breithaupt A, Schon J, Trujillo JD, Nooruzzaman M, Kwon T, Ahrens AK, Britzke T, McDowell CD, Piesche R, Singh G, Pinho Dos Reis V, Kafle S, Pohlmann A, Gaudreault NN, Corleis B, Ferreyra FM, Carossino M, Balasuriya UBR, Hensley L, Morozov I, Covaleda LM, Diel DG, Ulrich L, Hoffmann D, Beer M, Richt JA. 2025. H5N1 clade 2.3.4.4b dynamics in experimentally infected calves and cows. Nature 637:903–912.

37. Wernike K, Böttcher J, Amelung S, Albrecht K, Gartner T, Donat K, Beer M. 2022. Antibodies against SARS-CoV-2 Suggestive of Single Events of Spillover to Cattle, Germany. Emerg Infect Dis 28:1916–1918.

38. Michelitsch A, Dalmann A, Wernike K, Reimann I, Beer M. 2019. Seroprevalences of Newly Discovered Porcine Pestiviruses in German Pig Farms. Vet Sci 6.

39. Schulz D, Aebischer A, Wernike K, Beer M. 2024. No evidence of spread of Linda pestivirus in the wild boar population in Southern Germany. Virol J 21:205.

40. Linster M, Schrauwen EJA, van der Vliet S, Burke DF, Lexmond P, Bestebroer TM, Smith DJ, Herfst S, Koel BF, Fouchier RAM. 2019. The Molecular Basis for Antigenic Drift of Human A/H2N2 Influenza Viruses. J Virol 93.

41. Günther A, Wassermann J, Heck J, Bussi M, Aebischer A, Staubach C, Bergmann H, Leendertz FH, Beer M, Schares G, Wernike K. 2025. Opportunity Drives Spillover: Serological Surveillance across Carnivores, Omnivores and Herbivores in an HPAIV H5 Hotspot in North-East Germany, 2023–2025. bioRxiv doi:10.1101/2025.09.30.678011:2025.09.30.678011.

42. Daniel K, Ullrich L, Ruchnewitz D, Meijers M, Halwe NJ, Wild U, Eberhardt J, Schön J, Stumpf R, Schlotz M, Wunsch M, Girao Lessa L, Abd El-Whab E-SM, Kuryshko M, Dietrich C, Pinger A, Schumacher A-L, Germer M, Rohde M, Kukat C, Gieselmann L, Gruell H, Hoffmann D, Beer M, Erren T, Lässig M, Kreer C, Klein F. 2025. Detection of pre-existing humoral immunity against influenza virus H5N1 clade 2.3.4.4b in unexposed individuals. bioRxiv doi:10.1101/2025.01.22.634277:2025.01.22.634277.

43. Gauger PC, Vincent AL. 2020. Serum Virus Neutralization Assay for Detection and Quantitation of Serum Neutralizing Antibodies to Influenza A Virus in Swine. Methods Mol Biol 2123:321–333.

44. Kok A, Wilks SH, Tureli S, James SL, Bestebroer TM, Burke DF, Funk M, van der Vliet S, Spronken MI, Rijnink WF, Pattinson DJ, de Meulder D, Rosu ME, Lexmond P, van den Brand JMA, Herfst S, Smith DJ, Fouchier RAM, Richard M. 2025. A vaccine central in A(H5) influenza antigenic space confers broad immunity. Nature 647:1005–1013.

45. Stech J, Stech O, Herwig A, Altmeppen H, Hundt J, Gohrbandt S, Kreibich A, Weber S, Klenk HD, Mettenleiter TC. 2008. Rapid and reliable universal cloning of influenza A virus genes by target-primed plasmid amplification. Nucleic Acids Res 36:e139.

46. Burbelo PD, Ching KH, Bush ER, Han BL, Iadarola MJ. 2010. Antibody-profiling technologies for studying humoral responses to infectious agents. Expert Rev Vaccines 9:567–78.

47. Burbelo PD, Lebovitz EE, Notkins AL. 2015. Luciferase immunoprecipitation systems for measuring antibodies in autoimmune and infectious diseases. Transl Res 165:325–35.

48. Beigel JH, Voell J, Huang CY, Burbelo PD, Lane HC. 2009. Safety and immunogenicity of multiple and higher doses of an inactivated influenza A/H5N1 vaccine. J Infect Dis 200:501–9.

49. Watcharatanyatip K, Boonmoh S, Chaichoun K, Songserm T, Woratanti M, Dharakul T. 2010. Multispecies detection of antibodies to influenza A viruses by a double-antigen sandwich ELISA. J Virol Methods 163:238–43.

50. Rojas A, Germitsch N, Oren S, Sazmand A, Deak G. 2024. Wildlife parasitology: sample collection and processing, diagnostic constraints, and methodological challenges in terrestrial carnivores. Parasit Vectors 17:127.

51. Gardner IA, Hietala S, Boyce WM. 1996. Validity of using serological tests for diagnosis of diseases in wild animals. Rev Sci Tech 15:323–35.

52. Jensen TH, Andersen JH, Hjulsager CK, Chriel M, Bertelsen MF. 2017. Evaluation of a Commercial Competitive Enzyme-Linked Immunosorbent Assay for Detection of Avian Influenza Virus Subtype H5 Antibodies in Zoo Birds. J Zoo Wildl Med 48:882–885.

53. Lebarbenchon C, Brown JD, Luttrell MP, Stallknecht DE. 2012. Comparison of two commercial enzyme-linked immunosorbent assays for detection of Influenza A virus antibodies. J Vet Diagn Invest 24:161–5.

54. Lundqvist ML, Middleton DL, Radford C, Warr GW, Magor KE. 2006. Immunoglobulins of the non-galliform birds: antibody expression and repertoire in the duck. Dev Comp Immunol 30:93–100.

55. Magor KE. 2011. Immunoglobulin genetics and antibody responses to influenza in ducks. Dev Comp Immunol 35:1008–16.

56. de los Rios M, Criscitiello MF, Smider VV. 2015. Structural and genetic diversity in antibody repertoires from diverse species. Curr Opin Struct Biol 33:27–41.

57. Hawman DW, Tipih T, Hodge E, Stone ET, Warner N, McCarthy N, Granger B, Meade-White K, Leventhal S, Hatzakis K, Park S, Gaffney K, Rosenke K, Erasmus JH, Feldmann H. 2025. Clade 2.3.4.4b but not historical clade 1 HA replicating RNA vaccine protects against bovine H5N1 challenge in mice. Nat Commun 16:655.

58. Graaf A, Piesche R, Sehl-Ewert J, Grund C, Pohlmann A, Beer M, Harder T. 2023. Low Susceptibility of Pigs against Experimental Infection with HPAI Virus H5N1 Clade 2.3.4.4b. Emerg Infect Dis 29:1492–1495.

59. Kwon T, Trujillo JD, Carossino M, Machkovech HM, Cool K, Lyoo EL, Singh G, Kafle S, Elango S, Vediyappan G, Wei W, Minor N, Matias-Ferreyra FS, Morozov I, Gaudreault NN, Balasuriya UBR, Hensley LE, Diel DG, Ma W, Friedrich TC, Richt JA. 2025. Pathogenicity and transmissibility of bovine-derived HPAI H5N1 B3.13 virus in pigs. Emerg Microbes Infect 14:2509742.

60. Sanz-Muñoz I, Sánchez-Martínez J, Rodríguez-Crespo C, Concha-Santos CS, Hernández M, Rojo-Rello S, Domínguez-Gil M, Mostafa A, Martinez-Sobrido L, Eiros JM, Nogales A. 2025. Are we serologically prepared against an avian influenza pandemic and could seasonal flu vaccines help us? mBio 16:e0372124.

61. Corti D, Suguitan AL, Jr., Pinna D, Silacci C, Fernandez-Rodriguez BM, Vanzetta F, Santos C, Luke CJ, Torres-Velez FJ, Temperton NJ, Weiss RA, Sallusto F, Subbarao K, Lanzavecchia A. 2010. Heterosubtypic neutralizing antibodies are produced by individuals immunized with a seasonal influenza vaccine. J Clin Invest 120:1663–73.

62. Nachbagauer R, Wohlbold TJ, Hirsh A, Hai R, Sjursen H, Palese P, Cox RJ, Krammer F. 2014. Induction of broadly reactive anti-hemagglutinin stalk antibodies by an H5N1 vaccine in humans. J Virol 88:13260–8.

